# Restricted nucleation and piRNA-mediated establishment of heterochromatin during embryogenesis in *Drosophila miranda*

**DOI:** 10.1101/2021.02.16.431328

**Authors:** Kevin H.-C. Wei, Carolus Chan, Doris Bachtrog

## Abstract

Heterochromatin is a key architectural feature of eukaryotic genomes, crucial for silencing of repetitive elements and maintaining genome stability. Heterochromatin shows stereotypical enrichment patterns around centromeres and repetitive sequences, but the molecular details of how heterochromatin is established during embryogenesis are poorly understood. Here, we map the genome-wide distribution of H3K9me3-dependent heterochromatin in individual embryos of *D. miranda* at precisely staged developmental time points. We find that canonical H3K9me3 enrichment patterns are established early on before cellularization, and mature into stable and broad heterochromatin domains through development. Intriguingly, initial nucleation sites of H3K9me3 enrichment appear as early as embryonic stage3 (nuclear cycle 9) over transposable elements (TE) and progressively broaden, consistent with spreading to neighboring nucleosomes. The earliest nucleation sites are limited to specific regions of a small number of TE families and often appear over promoter regions, while late nucleation develops broadly across most TEs. Early nucleating TEs are highly targeted by maternal piRNAs and show early zygotic transcription, consistent with a model of co-transcriptional silencing of TEs by small RNAs. Interestingly, truncated TE insertions lacking nucleation sites show significantly reduced enrichment across development, suggesting that the underlying sequences play an important role in recruiting histone methyltransferases for heterochromatin establishment.

## Introduction

The separation of eukaryotic genomes into transcriptionally active euchromatin and silenced heterochromatin is a fundamental aspect of eukaryotic genomes (Allshire and Madhani 2018). Heterochromatin is the gene-poor, transposon-rich, late-replicating and tightly packaged chromatin compartment that was first cytologically defined over 90 years ago (Heitz 1928), in contrast to euchromatin, the gene-rich, lightly packed form of chromatin. These domains show characteristic distributions across eukaryotic genomes and are distinguished by unique sets of histone modifications (Peng and Karpen 2008; Elgin and Reuter 2013). Heterochromatin is found predominately at repetitive sequences, which mainly correspond to pericentromeres, the dot and the Y chromosome in flies, and is marked by tri-methylation of histone H3 lysine 9 (H3K9me3) (Elgin and Reuter 2013).

Heterochromatin formation, and the boundary between heterochromatic and euchromatic domains is established during early development. In *Drosophila melanogaster,* constitutive heterochromatin is not observed cytologically in the initial zygote, but emerges during blastoderm formation (Vlassova *et al*. 1991; Lu *et al*. 1998) (developmental stage 4; see **Figure 1A, B**). Chromatin assembly during this period is prior to any widespread zygotic transcription, and dependent on maternally loaded RNA and proteins (Elgin and Reuter 2013). Analysis of an inducible reporter gene has found that silencing occurs at the onset of gastrulation (end of stage 6), about 1 hour after heterochromatin is visible cytologically (Lu *et al*. 1998). The extent of silencing increases as embryonic development progresses, and by stage 15 (dorsal closure, between 11.5 and 13 hours of development), silencing patterns reminiscent of those for third instar larvae are established (Lu *et al*. 1998).

**Figure 1.**
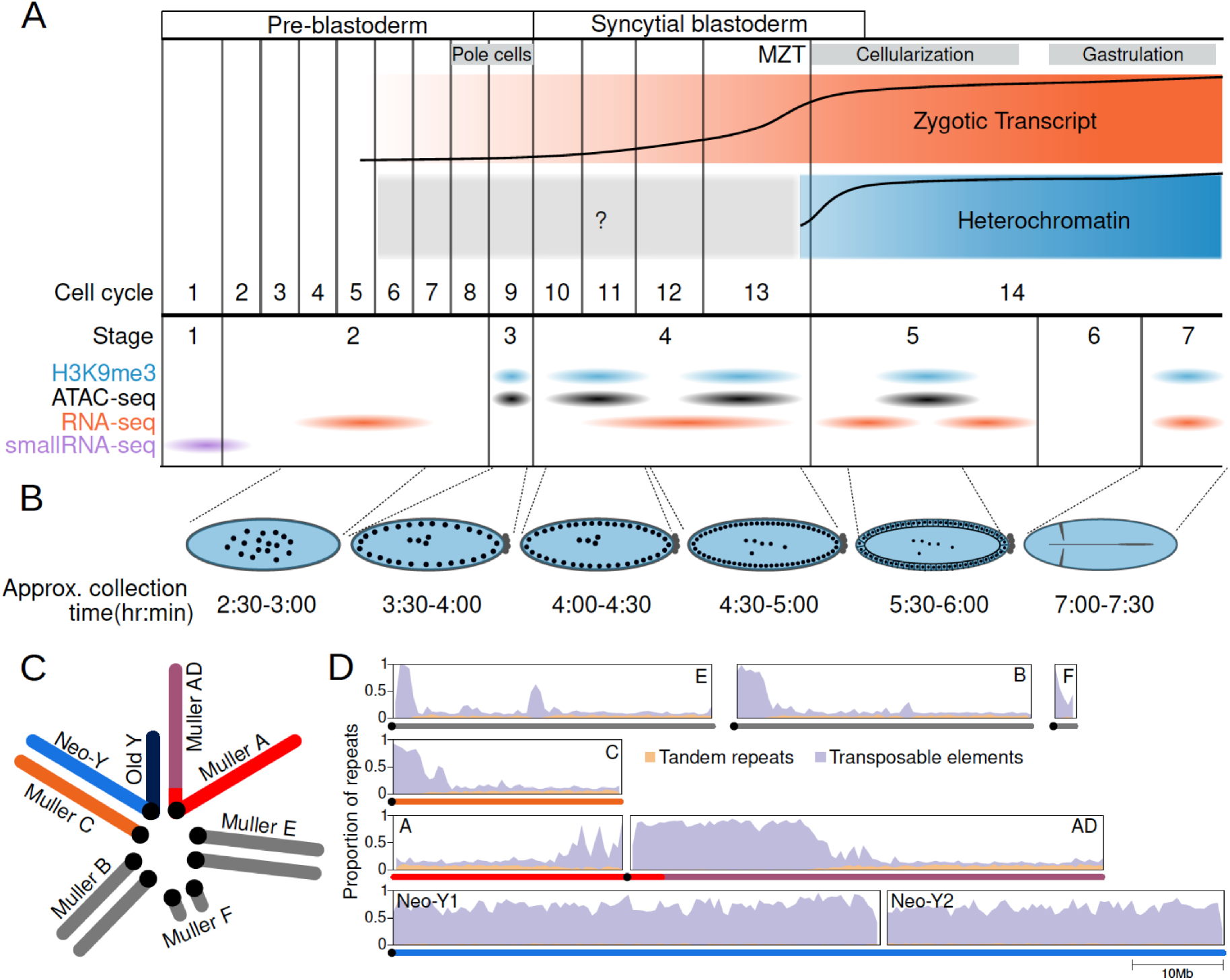
The repeat-rich genome of *Drosophila miranda* as a model to characterize heterochromatin establishment. A. Time course of embryonic development. Major land marks including the maternal and zygotic transition (MZT) are labeled. The orange and blue track depicts the approximate amount of zygotic expression and heterochromatin, respectively, throughout development. The cell cycle numbers and their corresponding embryonic stages are labeled. B. Cartoon diagram of the embryonic landmarks used for staging. The approximate time points of embryo collection are labeled under the embryo diagrams. C. Karyotype of *D. miranda* male. Muller elements are labeled and the sex chromosomes are color coded: neo-Y (blue), ancestral Y (dark navy), neo-X (orange), and X (red and purple). D. Repeat content of the *D. miranda* genome assembly. The cumulative repeat content for each chromosome is depicted, with the tandem repeats in orange and transposable elements in purple; unless otherwise stated, these repeat classes will be distinguished by these colors throughout the manuscript. Karyotypes are drawn below the graphs with the chromosomes color coded as in C.

The role of H3K9me3-associated heterochromatin in genome regulation and silencing of repetitive elements is well established. H3K9me3 is deposited by a conserved class of histone methyltransferases (Rea *et al*. 2000; Nakayama *et al*. 2001) and provides a high-affinity binding site for HP1 proteins (Bannister *et al*. 2001; Lachner *et al*. 2001). HP1 proteins can oligomerize, resulting in local chromatin compaction (Hiragami-Hamada *et al*. 2016), and help establish heterochromatin domains via phase separation (Larson *et al*. 2017; Strom *et al*. 2017). HP1 proteins can also recruit additional histone methyltransferases, thereby enabling spreading of heterochromatin (Canzio *et al*. 2013). Thus, H3K9me3 deposition is a crucial step in the formation of heterochromatin and the suppression of repetitive sequences, but the molecular mechanisms of H3K9me3 targeting and recruitment are not fully understood.

Several studies have suggested that small RNA-mediated silencing pathways can initiate the formation of heterochromatin (Holoch and Moazed 2015; Allshire and Madhani 2018). This was first identified in fission yeast, where mutations in components of the RNAi pathway showed disrupted heterochromatin silencing in centromeres (Volpe *et al*. 2002; Hall *et al*. 2002). In flies and mammals, a different class of small RNAs, PlWI-associated small RNAs (piRNAs; 23-29bps in Drosophila) are involved in the transcriptional silencing of TEs (Brennecke *et al*. 2007). piRNAs are derived from TE sequences and provide heritable libraries that target TE transcripts for degradation (Brennecke *et al*. 2007). Independent of the post-transcriptional cleavage activity, the piRNA pathway is required for transcriptional silencing of TE insertions in the ovary, as loss of PIWI causes the loss of heterochromatic repression through H3K9me3 (Sienski *et al*. 2012)(Wang and Elgin 2011; Darricarrère *et al*. 2013). Heterochromatin formation guided by piRNAs requires transcription of the target locus (co-translational silencing); Piwi-piRNA complexes bind to complementary, nascent TE transcripts and induce heterochromatin formation by recruiting heterochromatin factors, including histone methyltransferases for H3K9me3 deposition (Sienski *et al*. 2012) (Wang and Elgin 2011) (Le Thomas *et al*. 2013) (Yu *et al*. 2015; Batki *et al*. 2019).

In the developing embryo, maternally deposited PIWI is similarly required for the formation of H3K9me3 at the maternal zygotic transition during late blastoderm (Gu and Elgin 2013). However, the process and precise mechanism of rapid heterochromatin establishment across a large fraction of the genome such as the pericentromere and the Y chromosome remains poorly understood. Early cytological studies showed that heterochromatin becomes visible during blastoderm (Vlassova *et al*. 1991; Lu *et al*. 1998). More recently, *in vivo* immunofluorescence staining on the developing Drosophila embryo has shown that heterochromatin appears as early as cell cycle 13 (late stage 4) and rapidly increases through blastulation (cell cycle 14, stage 5) and requires the histone methyltransferase SetDB1 (Yuan and O’Farrell 2016; Seller *et al*. 2019). However, such imaging approaches lack resolution at genome-scale and are necessarily limited by the concentration and density of the targeted sequence and proteins. Emerging heterochromatin, therefore, may not yield strong enough signal for fluorescent detection. Hence, highly sensitive genomewide analyses of H3K9me3-dependent heterochromatin dynamics in early embryos are needed.

Here, we investigate the establishment of heterochromatin during early embryogenesis in *Drosophila miranda.* While this species lacks the genetic tools available in *D. melanogaster,* several features make *D. miranda* ideal for studying repetitive sequences and heterochromatin on a genome scale. *D. miranda* has a large repeat-rich neo-Y chromosome which formed through the fusion of an autosome (Muller-C) with the ancestral Y around 1.5 million years ago and has since more than doubled in size due to the accumulation of transposable elements. The high quality genome assembly has near end-to-end assemblies of most chromosome arms that include large fractions of heterochromatin (**Figure 1C-D,** (Mahajan *et al*. 2018)): roughly 30Mb of pericentromeric heterochromatin at the X and autosomes, and over 100Mb of the repeatrich Y chromosome have been assembled. In contrast, the most repeat-rich genome assembly of *D. melanogaster* contains only 14.6Mb of Y-linked sequence scattered across 106 contigs (Chang and Larracuente 2019). *D. miranda’s* neo-Y chromosome is predominantly assembled into two massive contigs (53.8 and 37.2 Mb) and harbors an abundance of active single-copy genes interspersed in repeats; the mixture of genes and repeats create nonredundant junctions facilitating sensitive read mapping (Wei *et al*. 2020). Additionally, unlike *D. melanogaster* which contains Mb-sized blocks of simple satellite DNA, this species has few simple satellites in its genome (Wei *et al*. 2018), thus allowing us to map most heterochromatic regions using ChIP-seq technology. We adapted an ultra-low input ChIP-seq protocol using single precisely staged embryos to investigate H3K9me3 profiles during early development (**Figure 1A**), to study the formation of H3K9me3-dependend heterochromatin. Our dense sampling during early embryogenesis (genome-wide H3K9me3 profiles for >70 individual embryos) allowed us to infer spatiotemporal heterogeneity in heterochromatin establishment, and evaluate the relationship between the nucleation and spreading of H3K9me3 marks to zygotic genome activation and maternally deposited piRNAs.

## Results

### Genome-wide profiling of H3K9me3 in Drosophila across an embryonic time course

To define the chromatin landscape before, during and after the establishment of heterochromatin, we collected *D. miranda* (MSH22) embryos from population cages at 18°C between 210-420 min, to target embryonic stage 3 (before the onset of heterochromatin formation), early- and late stage 4 (when some heterochromatin is first detected), stage 5 (when heterochromatin becomes cytologically visible), and stage 7 (when heterochromatin exerts its silencing effect), respectively (see **Figure 1A** and **Supplementary Table 1**). Note that embryonic development is highly stereotypical across the Drosophila genus, with nearly identical morphological landmarks (Kuntz and Eisen 2014). Embryos were examined under a light microscope to only select embryos of the correct stages, based on major morphological features to classify developmental stages (see Methods, **Figure 1B;** (Lott *et al*. 2014)). For each stage, we collected between 8 to 20 embryos each, which were subjected to single-embryo, ultra-low input ChIP-seq library preparation (Brind’Amour *et al*. 2015) with antibodies against the repressive histone mark H3K9me3. The large numbers of replicates (**Supplementary Table 1**) ensure robust and sensitive profiles of H3K9me3 enrichment across the stages. We initially performed two independent IP’s on individual embryos, targeting both H3K9me2 and H3K9me3 (**Supplementary Figure 1**). Overall, their enrichment profiles are very similar and highly correlated (**Supplementary Figure 1**). In addition, we spiked the libraries with stage 7 *D. melanogaster* embryos so that enrichment profiles can be compared across samples and stages after spike-in normalization (see Materials and methods and **Supplementary Figure 2**).

### Bulk of heterochromatin establishment occurs during early stage 4 of embryonic development

Based on cytology and microscopy, heterochromatin is first detectable at the last cell cycles in stage 4 (that is around nuclear cycle 12/13) and then becomes established rapidly during cellularization at stage 5 (Vlassova *et al*. 1991; Lu *et al*. 1998; Yuan and O’Farrell 2016; Larson *et al*. 2017; Strom *et al*. 2017). However, we already see clear H3K9me3 enrichment around the pericentric regions of each chromosome at early stage 4 (that is, nuclear cycle 10/11), followed by subsequent increases in enrichment through stage 7 (**Figure 2A**). The same pattern of establishment at early stage 4 is also apparent when we inferred regions of the genome with H3K9me3 enrichment peaks; the number of H3K9me3 peaks identified sharply increases from stage 3 (n = 2,202) to early stage 4 (n = 51,202), followed by proportionally smaller increases in peak numbers through the later stages (**Figure 2B** and **Supplementary Figure 3**). Interestingly, as development progresses, peak sizes broaden and become less sharp (**Figure 2B, C** and **Supplementary Figure 4**). After stage 3, we see a gradual increase of H3K9me3 enrichment around early peaks as embryos age (**Figure 2C**). Concordantly, regions of the genome where peaks are inferred in late stage 4, stage 5 and stage 7, already show positive H3K9me3 enrichment at early stage 4 on average (**Figure 2D**), even though they are depleted, on average, at stage 3 (**Figure 2D**). In fact, nearly 60% of the 116,791 peaks inferred for stage 7 are already enriched for H3K9me3 at early stage 4, even if the enrichment level is below the detection limit of peak calling (**Figure 2E**). Further, an additional 27% of the peaks also show enrichment at stage 3 (**Figure 2E**). Similarly, for late stage 4 and stage 5, the majority of the peaks are enriched in early stage 4. Altogether, these results indicate that pericentric heterochromatin establishment in *D. miranda* likely starts at or before stage 4, followed by increases in heterochromatin levels during subsequent stages.

**Figure 2.**
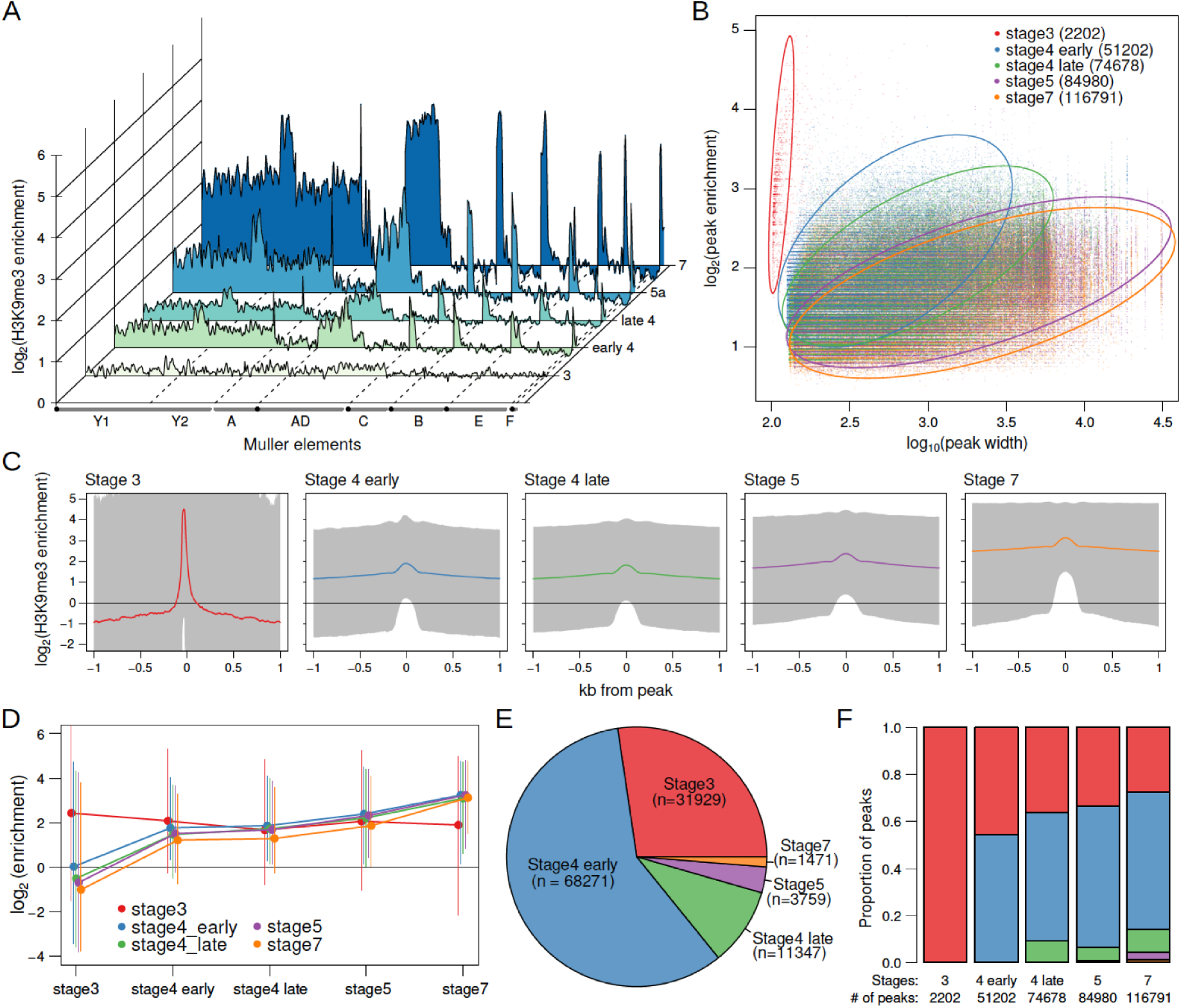
Developmental trajectory of heterochromatin enrichment and peaks. A. The genome-wide H3K9me3 enrichment landscape through five embryonic stages. The karyotypes are depicted below the X-axis, with the centromeres marked by black circles. B. The width and height of H3K9me3 peaks (points) as determined by MACS2, are plotted in log scale on the X- and Y-axes, respectively. The stages are color labeled with number of peaks in parentheses. Circles outline areas in which the bulk of the points of a stage resides. Unless otherwise stated, the developmental stages will be differentiated consistently with these colors henceforth. C. Median H3K9me enrichment in log scale is plotted ± 1kb around peaks for each stage. Gray area demarcates the 95% confidence intervals. D. H3K9me3 enrichment trajectory of peaks called for each stage across development. For every set of peaks called in each stage, the median enrichment value is plotted and connected across all developmental stages with the 95% CI demarcated by vertical lines. For example, Red points and lines are the enrichment values around stage3 peaks across all five stages. Points and CIs are horizontally staggered for clarity. E. Colored areas in pie chart mark the proportion of stage 7 peaks that are already enriched (>1.5-fold enrichment) in previous stages. F. Barplot format of E, but for peaks called in every stage.

### Bona fide stage 3 peaks nucleate heterochromatin

While the bulk of H3K9me3 enrichment appears at early stage 4, we also identified a small number of peaks at stage 3 (n = 2,202) that showed sharp and narrow enrichment around their center that precipitously drops to depletion less than 100bp away (**Figures 2B, C**). This is a strikingly different pattern from peaks called at all other stages which show elevated enrichment that span well over 1kb away from the peak (**Figure 2B, C**). To ensure that stage 3 peaks and enrichment are not artifacts due to the small number of nuclei (‘phantom peaks’), we collected ChIP-seq of stage 3 embryos using three additional antibodies targeting H3, H3K4me3 (an active transcription mark), and H4K16ac (the dosage compensation mark). We find that many of the stage 3 heterochromatin peaks show similarly elevated enrichment in the other preparations, indicating that a subset of the peaks are indeed artifacts (Jain *et al*. 2015) (**Figure 3A**). We therefore removed 1205 peaks which show enrichment in one or more of the other ChIP preparations, resulting in a set of 997 H3K9me3-specific peaks (henceforth stage 3 peaks; **Figure 3A and Supplementary figure 5**).

**Figure 3.**
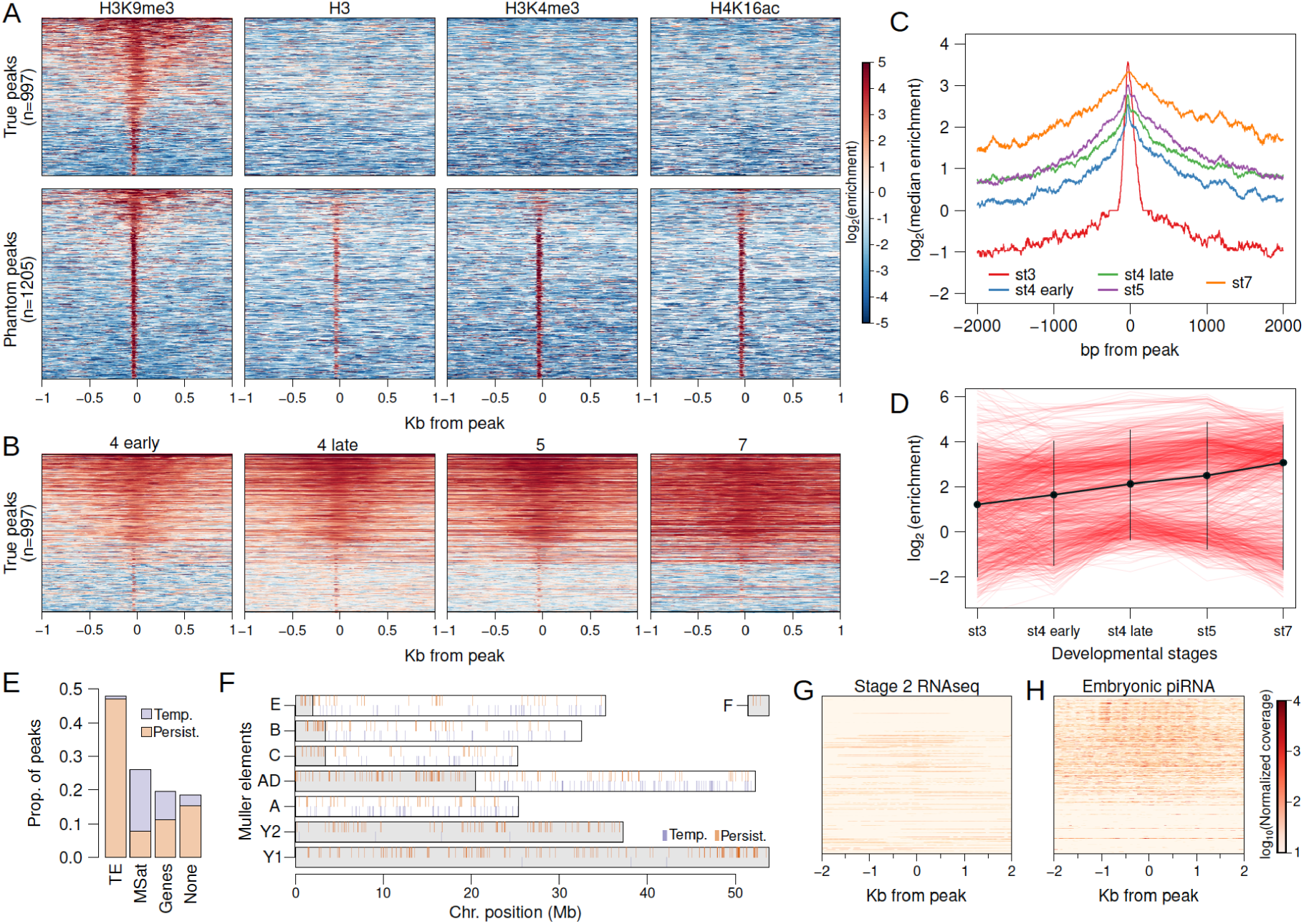
Stage 3 H3K9me3 peaks nucleate heterochromatin. A. Separation of genuine stage 3 peaks from nonspecific (phantom) stage 3 peaks with ChIP-seqs against alternative histone modifications. Heatmaps depict the extent of enrichment around stage3 H3K9me3 peaks (±1kb). Peaks are sorted by the average H3K9me3 enrichment around each peak, therefore; all the different ChIP-seqs have the same ordering. Top panels are H3K9me3-specific peaks, while bottom panels are non-specific peaks. B. H3K9me3 enrichment around stage3 peaks across developmental stages; peaks are ordered as in A (top panel). C. Median H3K9me3 enrichment across developmental stages around stage3 peaks. D. Developmental trajectory of each H3K9me3 peak (red lines). Enrichment of a peak is estimated as enrichment averaged across ±100bp around each peak. The average across all peaks is in black with error bars representing 95% confidence intervals. E. Distribution of stage3 peaks across different annotation categories. F. Placement of peaks across the genome; gray regions mark the pericentric and heterochromatic regions of the genome. G,H. Maternal deposited RNA and piRNAs around (±1kb) stage 3 peaks; peaks are ordered as in A (top panel).

Around stage 3 peaks, H3K9me3 enrichment gradually expands across development (**Figure 3B, C**), emblematic of the spreading of the heterochromatic domain. Since a single nucleosome is wrapped by ~147bp of DNA, the narrow width of the stage 3 peaks (median width of 125bp) suggests that they predominantly represent single nucleosomes that nucleate the deposition of H3K9me3 in neighboring nucleosomes in subsequent stages (**Figure 3A, C**). 52.5% and 28.7% of the stage 3 peaks overlap with annotated TEs and microsatellites (**Figure 3E**). Unexpectedly, though, they are found scattered across the genome, as opposed to being concentrated near the pericentric regions (**Figure 3F**). Interestingly, 25.6% (n=256) of the stage3 peaks appear to become depleted of H3K9me3 enrichment over time (**Figure 3B, D**), possibly indicating that some nucleating sites fail to initiate and/or maintain heterochromatin spreading, but they could also be phantom peaks. The bulk of these temporary peaks overlap microsatellites (71.4%) and only 3% overlap TEs (**Figure 3E**); in contrast, peaks that persist and expand through development (n=741) mostly overlap TEs (63.3%), and only 10.5% are found at microsatellites (**Figure 3E**). Nearly all the temporary peaks are found outside of the pericentromere and the heterochromatic Y (**Figure 3F**).

H3K9me3 nucleation may be driven by RNA-mediated targeting (Gu and Elgin 2013). We utilized embryonic RNA-seq data (Lott *et al*. 2014) and sequenced the piRNAs in 0-1hr embryos to test for an association between H3K9me3 targeting and silencing of maternal or nascent TE transcripts by RNAi. We find very few RNA-seq reads from stage 2 or stage 4 embryos mapping to or around H3K9me3 peaks at embryonic stage 3 or 4 (**Figure 3G, Supplementary Figure 6**). Furthermore, while piRNA reads map abundantly around these peaks, they are not particularly enriched at or in close proximity to these peaks (**Figure 3H**). We note, though, that the temporary stage3 peaks are less associated with maternally deposited piRNAs (**Figure 3H**). While this may simply reflect that temporary peaks are artifacts or overlap few TEs (**Figure 3E**), it could also suggest that proper establishment and subsequent spreading of heterochromatin requires piRNA.

### Nucleation and spreading at early stage 4 are primarily at TEs and pericentric regions

While only about ~1000 H3K9me3 peaks are identified at stage 3, this number drastically increases to 51,202 in the early stage 4 samples (**Figure 4A**). Early stage 4 peaks show a gradual increase in levels of H3K9me3 enrichment and a local expansion of the H3K9me3 domain across development (**Figure 4A** and **Supplementary Figure 7**). Unlike stage 3 peaks, nearly all early stage 4 peaks (97.7%) remain enriched throughout development. To better characterize this rapid burst of nucleation activity, we divided the early stage 4 peaks into those that already show enrichment at stage 3 and those that do not, resulting in 16,630 and 35,977 peaks, respectively. The former (hence forth, old peaks) contain sites that are in the process of nucleation at stage 3 but their H3K9me3 enrichment is below the threshold used for peak-calling. The latter (hence forth, new peaks) contain sites that began nucleating at stage 4, as they show no prior heterochromatin enrichment.

**Figure 4.**
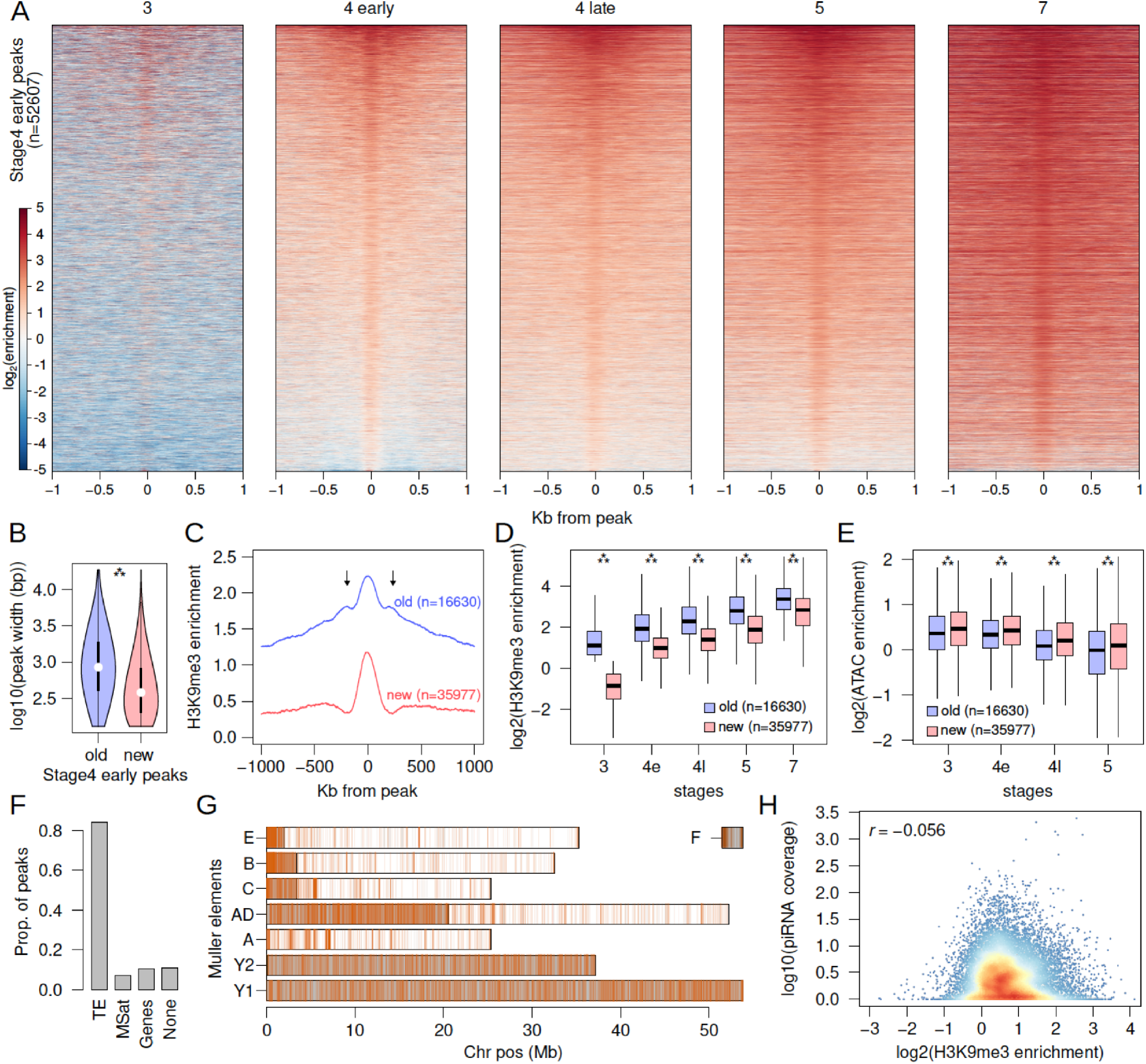
Rapid nucleation in early stage 4. A. H3K9me3 enrichment around (±1kb) early stage 4 peaks across development. Peaks are sorted by mean enrichment across early stage 4. B. Peak widths of stage4 peaks that show enrichment in stage 3 (old) and peaks that show no enrichment in stage (new); *** = p < 2.2e-16 (Wilcoxon Rank Sum Test). C. Median H3K9me3 enrichment around the old and new stage 4 peaks. Arrows mark secondary peaks around the old peaks. D. H3K9me3 enrichment of old and new stage 4 peaks across development; *** = p < 2.2e-16 (Wilcoxon Rank Sum Test). E. Accessibility as measured by ATAC-seq enrichment in old and new peaks; ** = p < 2.2e-16 (Wilcoxon Rank Sum Test). F. Distribution of new stage4 peaks across different annotation categories. G. Genome-wide distribution of new stage 4 peaks; gray regions mark the pericentric and heterochromatic regions of the genome. H. Density scatter plot of the correlation between H3K9me3 enrichment of new stage 4 peaks against maternally deposited piRNA coverage (*r* = Pearson’s correlation coefficient).

Consistent with subsequent spreading of H3K9me3 after nucleation, old peaks are on average 847 bp, significantly wider than young peaks which are on average 378 bp (Wilcoxon Rank Sum Test, two-tailed p-value < 2.2e-16; **Figure 4B**). Further, secondary peaks can be observed around the old peaks, indicating the deposition of H3K9me3 to adjacent nucleosomes by histone methyltransferases (**Figure 4C**). In contrast, the new peaks show a single summit flanked by depletion, consistent with H3K9me3 at single nucleosomes. Both sets of peaks increase in H3K9me3 enrichment across developmental stages with the new peaks having significantly lower enrichment across all stages; however, the difference in enrichment between the two sets of peaks decreases over time (**Figure 4D**). The gradual increase in H3K9me3 enrichment across stages likely reflects increasing numbers of nuclei/cells in the embryo with H3K9me3 deposited around nucleating sites as heterochromatin becomes stably established, with older peaks likely reaching saturation as the new peaks catch up. To determine whether increases in H3K9me3 enrichment are associated with expected decreases in chromatin accessibility, we generated ATAC-seq libraries from single embryos for the same developmental stages (absent stage 7). Indeed, we find that both sets of peaks show decreasing accessibility over development, with the old peaks being significantly less accessible across all stages (**Figure 4E**).

Expectedly, most of the stage 4 peaks (84.2%) overlap with annotated TEs (**Figure 4F**) and 82.6% of them are concentrated at the pericentromere and the repeat-rich Y chromosome (**Figure 4G**). These distributions are drastically different from stage 3 peaks, revealing that nucleation at stage 4 is more localized to canonically heterochromatic sites and transposable elements. Again, we looked for associations of stage 4 peaks with transcripts and piRNAs, but found no evidence that the peaks are enriched with either. As with the stage 3 peaks, very few stage 2 or 4 embryonic RNAseq reads map to or around the peaks (**Supplementary figure 8**). Further, piRNA reads are not significantly enriched at the peaks (**Supplementary figure 9**), and H3K9me3 enrichment at the peaks shows no correlation with the abundance of piRNA reads mapping around them (**Figure 4H**).

### Heterochromatin nucleation at TEs is restricted at stage 3 but wide at stage 4

As TEs dominate the stage 3 and the early stage 4 nucleation sites (**Figure 3E** and **4F**), we further characterized the pattern of nucleation and spreading at specific TE families annotated across the genome. Of the 235 entries in the TE library (Hill and Betancourt 2018), only 35 (14.8%) overlap with stage 3 peaks while 189 (80.3%) overlap with the early stage 4 peaks. Most of the stage 3 peaks reside in variants of the retrotransposons LOA (non-LTR), BEL (LTR), Gypsy11 (LTR), TRAM (LTR), and R1 (non-LTR) (**Figure 5A**). The early stage 4 peaks, on the other hand, show a very different distribution of peaks among TEs. For example, three TE variants (CR1-1, LOA-3 and Gypsy18) have the most early stage 4 peaks but few stage 3 peaks (**Figure 5A**), suggesting that some TEs nucleate heterochromatin earlier than others.

**Figure 5.**
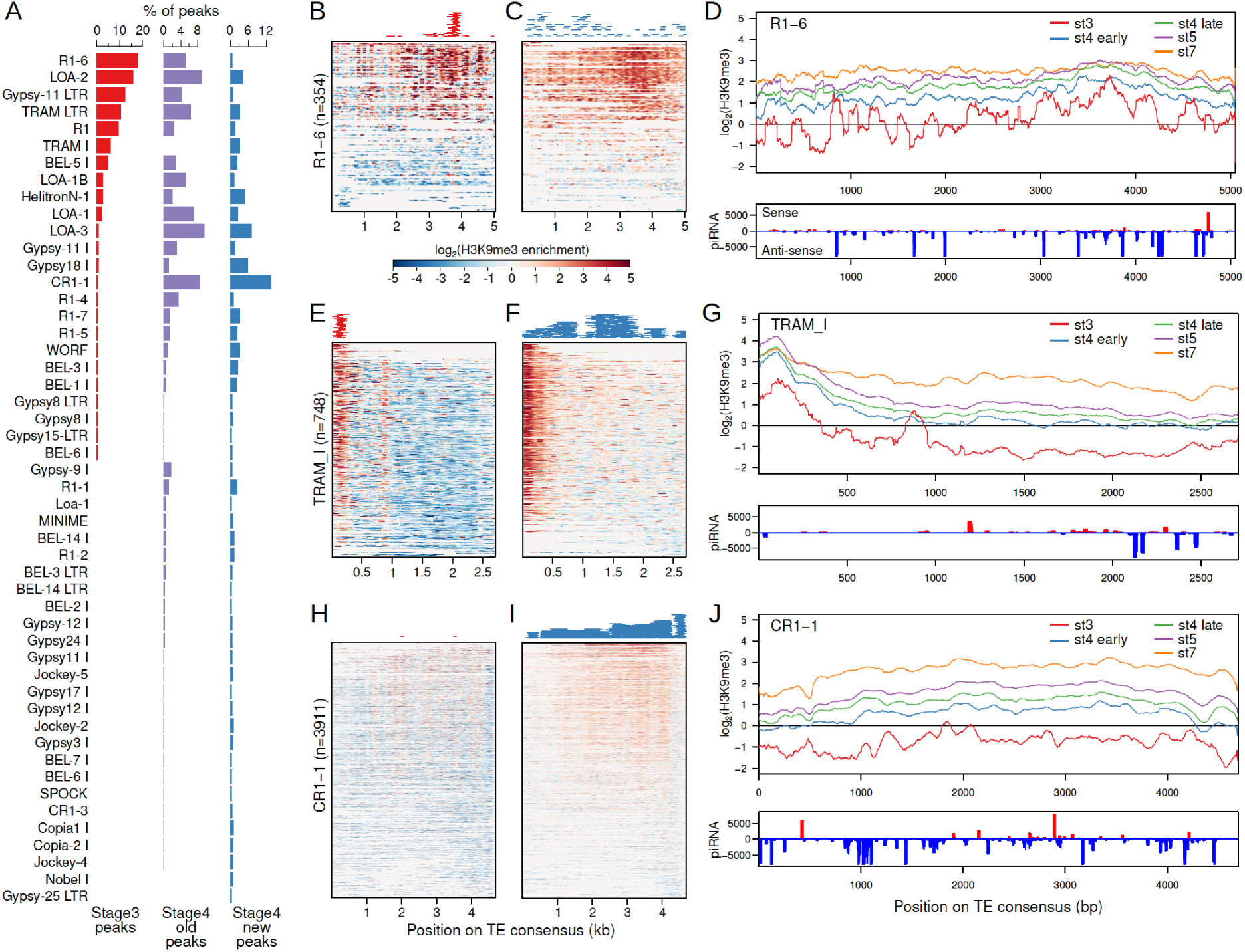
Narrow nucleation followed by wide establishment of heterochromatin at TEs. A. TEs with annotated insertions overlapping stage 3, stage 4 old peaks, and stage 4 new peaks are listed; barplots depict the proportion of peaks overlapping with each TE. B. H3K9me3 enrichment at stage 3 at all annotated TE insertions of the TE R1 variant (R1-6). Full length and fragmented annotated insertions are lined up with respect to their positions on the consensus TE sequences. Insertions are sorted by average enrichment. Positions of the called peaks are plotted above the heatmap. Same with B, but for stage 4 enrichment and peaks. D. Top. Mean enrichment of all insertions for the TE across development. Bottom. Sense and anti-sense piRNA mapping across the TE. E-G and H-J, same as B-C but for the TEs TRAM and CR1-1, respectively. For more examples of enrichment over TEs, see supplementary figure 10.

H3K9me3 nucleation within TEs could occur at specific positions. We looked at the placement of H3K9me3 peaks with respect to the full-length TE consensus sequence and overall H3K9me3 enrichment of all the annotated insertions for each TE. For TEs with large numbers of stage 3 peaks, the peaks are typically highly restricted to a specific region of TEs. For example, the vast majority of the stage 3 peaks localize between 3500-4000bp of the R1-6 element (**Figure 5B**). This starkly contrasts from the early stage 4 peaks, which are scattered across the entirety of the R1-6 element (**Figure 5C**). As expected, the position of the cluster of stage 3 peaks also corresponds to the region with the highest H3K9me3 enrichment across the R1-6 element, producing spikey and heterogeneous H3K9me3 enrichment profiles at stage 3 (**Figure 5B-D**). However, at later stages H3K9me3 enrichment becomes more evenly elevated around the peak region, indicative of spreading of heterochromatin for robust silencing during development (**Figure 5D**). Similarly, the TRAM element shows highly restricted H3K9me3 enrichment and peaks localized to the 5’ (0-300bp) region at stage 3 followed by broad heterochromatin enrichment at early stage 4 and peaks spread throughout the element (**Figure 5E-G**). The 5’ region of TRAM, where the early nucleating peaks reside, continues to increase in H3K9me3 over development as enrichment levels at the rest of the TE (**Figure 5G**). This pattern is also observed in other TEs with abundant stage 3 peaks (**Supplementary figure 10A,B**). Notably, the 5’ region of nearly all of the 748 TRAM copies found in the *D. miranda* genome show elevated H3K9me3 enrichment even though only 27 peaks have been called there (**Figure 5E**), indicating that peak calling is highly conservative and substantially underestimates the number of nucleation sites.

For TEs like CR1-1 that have few to no stage 3 peaks, H3K9me3 establishment at early stage 4 is nonetheless very broad (**Figure 5H-J, Supplementary figure 10C, D**). When looking at H3K9me3 enrichment at stage 3 across all CR1-1 insertions, we noticed low but consistent patterns of localized H3K9me3 enrichment at multiple positions of the TE (e.g. ~1800bp, ~2200bp, and ~4000bp); these places also appear to be some of the most elevated regions at later stages (**Figure 5J, Supplementary figure S7C, D**). These patterns suggest that despite the absence of H3K9me3 peak calls at stage 3, localized nucleation likely is in progress at this stage, but possibly in only a subset of nuclei, causing low overall enrichment. Thus, we conclude that nucleation begins first as targeted H3K9me3 enrichment at TE insertions followed by wide H3K9me3 deposition across the rest of the element.

### Early zygotic transcription and high maternal piRNA abundance associated with early nucleation

The piRNA pathway allows for sequence-specific targeting of TEs for both co-transcriptional silencing and post-transcriptional degradation (Malone and Hannon 2009). To determine whether piRNAs play a role in the targeted nucleation at TEs, we looked at the amount of maternally deposited sense and anti-sense piRNA mapping to TEs. Interestingly, we find that TEs with stage 3 peaks are significantly overrepresented with TEs generating a high abundance of sense and anti-sense piRNAs (**Figure 6A**; p = 4.2e-06 and p=0.000118, respectively, Wilcoxon Rank Sum Test).

**Figure 6.**
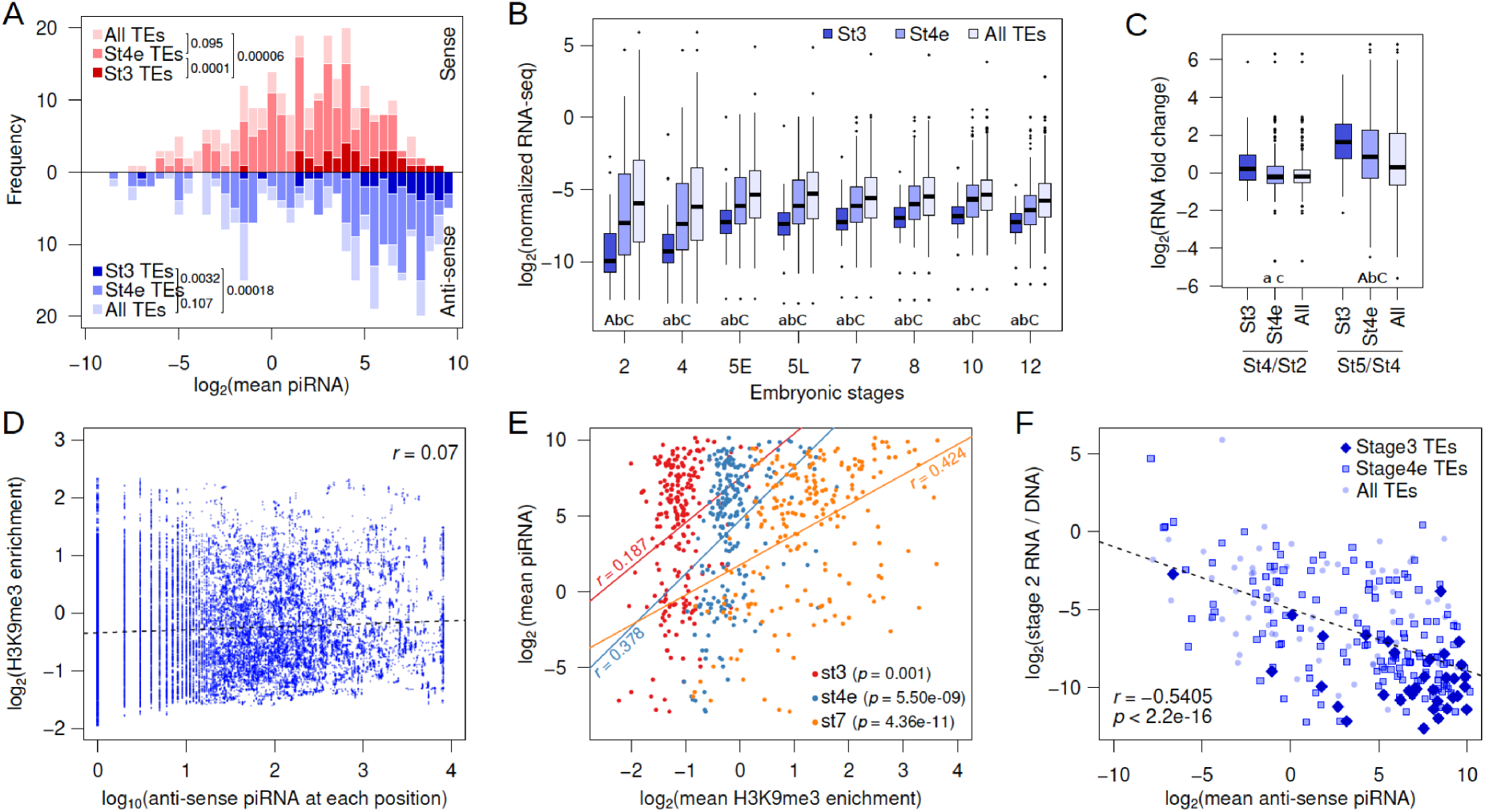
Association between early H3K9me3 nucleation at TEs, piRNA production, and expression. A. Distribution of TEs by their average maternally deposited sense (top, red) and anti-sense (bottom, blue) piRNA coverage. TEs with stage 3 peaks (stage 3 TEs), stage4 early peaks (stage4e TEs), and all TEs are in decreasing color intensity. P-values of pairwise comparisons determined by Wilcoxon’s Rank Sum Test are marked beside the legends. B. Difference between TE expression of TEs with stage 3 peaks, stage 4 peaks and all TEs across embryonic development. Pairwise significance is determined by Wilcoxon’s Rank Sum Test and represented by letters where the lower-case letters denote p-value < 0.05 and upper-case letters denote p-value < 0.001 after multiple-testing correction with false discovery rate; A = stage3 TEs vs. stage4 TEs, B = stage 4 early TEs vs. all TEs, and C = stage 3 TEs vs. all TEs. The absence of a letter denotes non-significance. C. Zygotic expression of different TE classes is approximated by the fold difference between early embryonic stages. D. For each position of stage 3 nucleating TEs, the H3K9me3 enrichment is plotted against the number of piRNA reads mapped. Linear regression is plotted in dotted line and Pearson’s correlation coefficient (*r*) is labeled. E. Correlation between average H3K9me3 enrichment across a TE at different developmental stages and the average piRNA abundance. Pearson’s correlation coefficients (*r*) are labeled beside the regression lines. F. Negative correlation between maternally deposited antisense piRNA and expression of all TEs (light blue dots), stage 3 TEs (dark blue diamonds), and stage 4 TEs (blue boxes). Expression of TEs is scaled by the copy number by dividing the RNA-seq read counts with DNA-seq read counts. Dotted line demarcates the linear regression and *r* represents the Pearsons’ correlation coefficient.

Co-transcriptional silencing of repeats requires early zygotic transcription of TEs. We examined TE expression in developmentally staged (stages 2 through 12) and sexed single embryo RNA-seq datasets (Lott *et al*. 2014); note that while these data measure standing levels of transcripts, changes in RNA transcript levels across stages reflect zygotic transcription, or RNA degradation. Across all developmental stages, transcript abundances of TEs with stage 3 peaks are significantly lower than the rest (**Figure 6B**). The vast majority of transcripts in stage 2 embryos are maternally deposited, and TE expression increases across the board as zygotic expression ramps up (most noticeable between stage 4 and 5). However, TEs with stage 3 nucleation sites show the highest and earliest increase in transcript abundance (**Figure 6B**). While transcript abundances of most TEs, on average, remain unchanged between stages 2 and 4 (0.97-fold difference), stage 3 nucleating TEs show a 1.41-fold increase in RNA abundance from stage 2 to stage 4 embryos (**Figure 6C**). Interestingly, TEs with stage 4 nucleation peaks, although showing no change in expression between stage 2 and 4 (1.01-fold difference), start showing evidence of zygotic transcription slightly later, when contrasting transcript abundances between zygotic stages 4 and 5 (**Figure 6B,C**). Theseresults reveal that early nucleating TEs are among the first to be zygotically transcribed during embryogenesis, consistent with the need for nascent TE transcripts to induce PIWI-associated co-transcriptional silencing (Shimada *et al*. 2016).

However, regions with abundant piRNA mapping on TEs do not correspond to the positions of stage 3 peaks nor the regions of elevated H3K9me3 enrichment (**Figure 5D,G,J**). Furthermore, along TEs with stage 3 peaks, the amount of piRNA mapping shows little-to-no correlation with H3K9me3 enrichment across the TE (**Figure 6D**). These results suggest that despite the correlation of TEs with stage 3 H3K9me3 peaks having a high abundance of maternally deposited piRNAs and showing early zygotic expression, piRNAs do not seem to provide sequence specificity for targeted nucleation of heterochromatin. Consistent with this, piRNA abundance is much more strongly and significantly correlated with H3K9me3 enrichment of TEs at the later developmental stages (**Figure 6C**) suggesting that piRNAs facilitate the spreading and maturation of H3K9me3-dependent silencing after early nucleation.

### Loss of nucleating sites at 5’ LTR diminishes heterochromatin establishment

Some TE’s, such as TRAM, have most of their stage 3 peaks and H3K9me3 enrichment in their 5’ region. This, along with early zygotic expression, raises the possibility that transcription initiation may be involved in nucleation. TRAM, a LTR retrotransposon, is flanked by two 372bp long terminal repeats (Steinemann and Steinemann 1997) (**Supplementary figure 11A**), which contains numerous stage 3 peaks (**Figure 5A**). For LTR retrotransposons, the 5’ LTR contains the primary promoter (Thompson *et al*. 2016). Indeed, the 5’ LTR of TRAM is highly enriched for H3K9me3 (**Figure 7A**, **Supplementary figure 11A**) and, to a lesser extent, also the 3’ LTR (**Figure 6A, Supplementary figure 11A**); we suspect the lower 3’ enrichment mostly reflects unavoidable non-unique mapping between identical sequences within the LTR. In addition, LTR retrotransposons can form head-to-tail tandems such that an 3’ LTR is also the 5’ LTR of a downstream element (McGurk and Barbash 2018; Ke and Voytas 1997), a structure which we indeed find in the *D. miranda* genome (**Supplementary figure 11B**). Other LTR retrotransposons show a similar enrichment of stage 3 peaks in their 5’ LTR (i.e. Gypsy-11, **Supplementary figure 12**).

**Figure 7.**
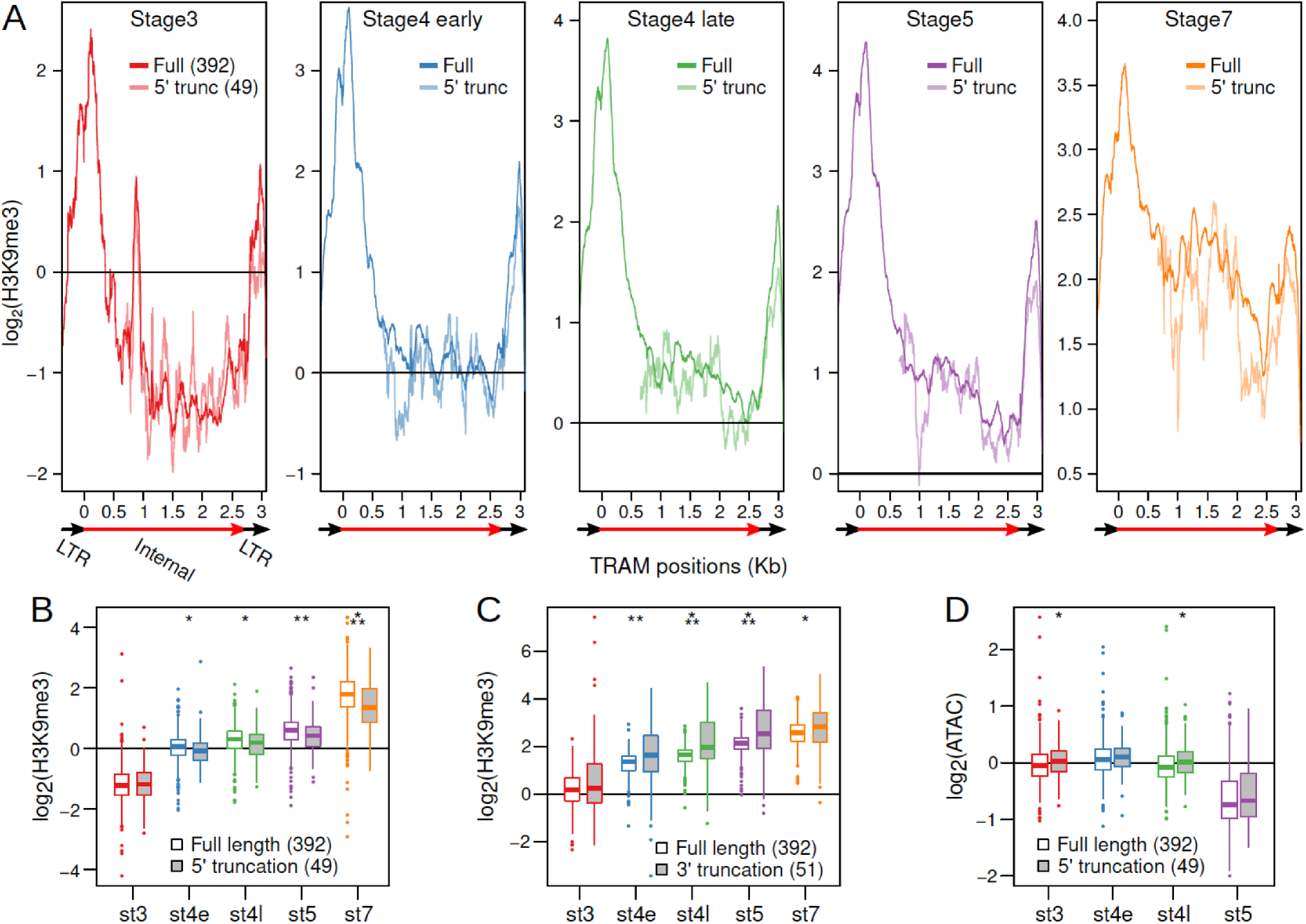
Loss of 5’ LTR and nucleation sites reduce H3K9me3 enrichment at TRAM insertions. A. H3K9me3 enrichment averaged across full length TRAM insertions (sold lines) and 5’ truncated TRAM insertions lacking the 5’ nucleation sites (dotted lines) in different developmental stages. Structure of TRAM is labeled below the plots (also see Supplementary figure 11). Note that the Y-axes change across the plots. B. Distribution of average H3K9me3 enrichment for full length (boxes with solid colors) and 5’ truncated (boxes with colored outlines) insertions in different developmental stages. For each insertion, enrichment is averaged across the last 1kb of the internal sequence (3’ LTR is excluded). Boxplots depict the distribution of H3K9me3 enrichment across insertions. C. Same as B, but averaged across the first 1kb of the internal sequence of full length and 3’ truncated insertions (5’ LTR is excluded). D. Same as B, but with ATAC enrichment instead of H3K9me3 enrichment. * = p < 0.05, ** = p < 0.005, *** = p < 0.00005, Wilcoxon’s Rank Sum test.

To determine whether the loss of 5’ LTR may perturb the establishment of heterochromatin, we identified TRAM insertions with 5’ truncations (n=49) abolishing the 5’ LTR and nucleating positions, and compared their H3K9me3 enrichment across development to that of full-length insertions (n=392) (**Figure 7A**). We find that 5’ truncated copies are significantly less enriched for H3K9me3 at their homologous regions after stage 3 with the difference in enrichment between 5’ truncated and full-length copies increasing throughout development (**Figure 7A,B**); at stage 7, full length inserts are on average 1.36-fold more enriched than 5’ truncated inserts. We note that the difference in enrichment is likely to be a severe underestimate, since non-unique cross-mapping (which is inevitable for repeats) between the different copies curtails the difference in read coverage and subsequently enrichment estimates between copies; the difference in enrichment therefore relies on reads that gap insertion junctions which are unique for different insertions. Additionally, we expect heterochromatic spreading from neighboring repeats to further minimize the difference in enrichment.

Notably, diminished H3K9me3 enrichment is not observed from 3’ truncated TRAM copies (n=51) where the 5’ LTR remains intact (**Figure 7C, Supplementary figure 13**); in fact, 3’ truncated TRAM copies have elevated enrichment compared to full length copies as they, by necessity, contain proportionally more of the highly enriched 5’ sequences (**Supplementary figure 13**). Further supporting the functional importance of the promoter adjacent nucleating sites for heterochromatin establishment, we find that 5’ truncated insertions of TRAM tend to be more accessible based on ATAC-seq enrichment; albeit, the differences are either marginally significant or insignificant (**Figure 7D**).

### Early nucleating TEs are robustly silenced but historically active elements

The amount of piRNA mapping to TEs is significantly anti-correlated with the maternally deposited transcripts abundance, particularly when scaled by TE copy number (Pearson’s correlation coefficient = – 0.54, p < 2.2e-16; **Figure 6F**). TEs with stage 3 nucleation are associated with high piRNAs counts (**Figure 6A**), and as expected, significantly overrepresented with lowly expressing TEs (**Figure 6A**; p = 1.582e-08, Wilcoxon’s Rank Sum Test); they remain to be lowly expressed as zygotic transcription increases after stage 4, indicating that early nucleating TEs are among the most repressed TEs (**Figure 6A**). TEs with stage 4 peaks show intermediate patterns with regards to piRNA counts and expression (**Figure 6A,B**).

High piRNA production and H3K9me3 enrichment and low transcript abundance of the early nucleating TEs suggest that they are targeted for robust silencing. These TEs may have high transposition rates and impose a high fitness burden, creating strong selective pressure to evolve and strengthen epigenetic silencing mechanisms. Indeed, we find that the early nucleating TEs are significantly more abundant in both the male and female *D. miranda* genomes (**Figure 8A**). Because genome-wide TE abundance can reflect ancient accumulation and not just recent transposition rates, we also looked at insertions specifically on the recently formed neo-Y chromosome. Consistent with recent activity of these TEs, the neo-Y has on average 1.79-fold more early nucleating TEs than the rest (**Figure 8B**). These results reveal that the early nucleating TEs have been recently active in the *D. miranda* lineage, generating large number of insertions throughout the genome.

**Figure 8.**
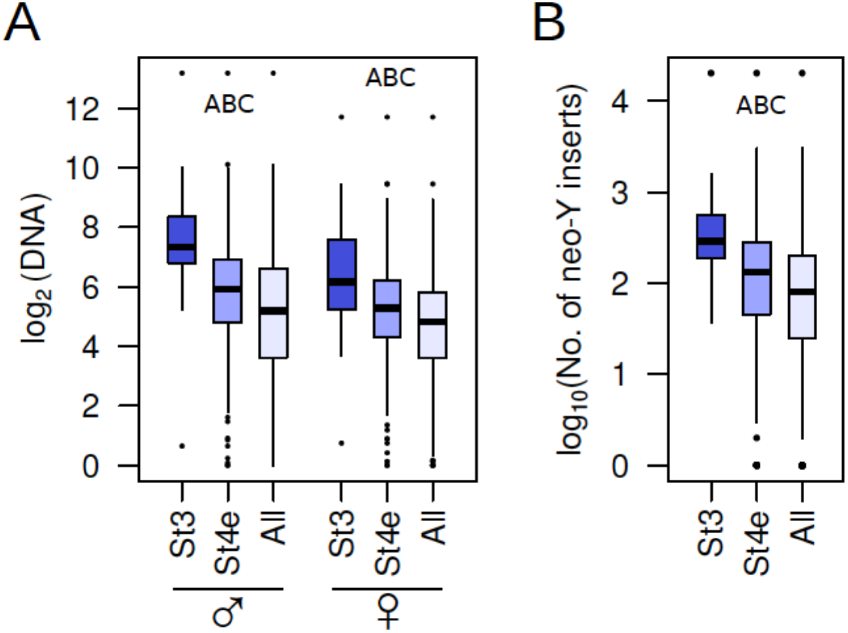
Genomic abundance of early nucleating TEs. A. Comparisons between the copy number abundance of stage3 nucleating, stage 4 nucleating, and all TEs in males and females. B. Same as A, but with number of insertions on the neo-Y. Pairwise significance is represented by letters where the lower-case letters denote p-value < 0.05 and upper-case letters denote p-value < 0.001 (Wilcoxon Rank Sum Test); A = stage3 TEs vs. stage4 TEs, B = stage 4 early TEs vs. all TEs, and C = stage 3 TEs vs. all TEs.

## Discussion

### H3K9me3-dependent heterochromatin nucleation in early development

In metazoans, extensive epigenetic reprogramming occurs during gametogenesis and early embryogenesis where most chromatin marks, including H3K9me3, are erased and re-established later (Wang *et al*. 2018). H3K9me3 re-establishment in somatic tissues is crucial for normal development in Drosophila and mammals. A number of different pathways have been identified that recruit H3K9 methyltransferases to target loci, and include a diversity of guides such as DNA binding proteins and non-coding RNAs. For example, in yeast small interfering RNAs (siRNAs) matching pericentric repeats are required for recruitment of the silencing machinery to pericentric heterochromatin (Volpe *et al*. 2002; Hall *et al*. 2002). In mammals, a parallel pathway exists based on DNA sequence-based recognition and silencing of mobile elements by KRAB-zinc-finger DNA-binding proteins. These proteins bind DNA via an array of zinc-fingers and recruit the universal co-repressor KAP1 (KRAB associated protein 1) through their KRAB domain. In turn, KAP1 recruits histonemethyltransferases to TE insertions to promote deposition of repressive chromatin marks (Wolf *et al*. 2015).

The role of H3K9me3 in early somatic development has been studied in mouse, and H3K9me3-dependent heterochromatin was shown to undergo dramatic reprogramming during early embryonic development (Wang *et al*. 2018). Targeting of H3K9me3 during fly embryogenesis is poorly understood. We find that establishment of heterochromatin begins with limited H3K9me3 deposition at single nucleosomes at or before stage 3. These nucleating sites expand gradually by H3K9me3 deposition to neighboring nucleosomes in subsequent stages to form mature heterochromatin. Unexpectedly, we find that early nucleation sites may not be restricted to the pericentric regions, and can be lost through development. Nevertheless, the majority of nucleating sites are localized to specific regions of transposable elements such that across different insertions, the same positions show elevated H3K9me3 enrichment. Interestingly, truncated insertions that lost these positions show significantly reduced H3K9me3 enrichment across the remainder of the TE and maintain elevated accessibility across development, revealing that these nucleating positions are important for subsequent local spreading of heterochromatin.

### Potential mechanisms for targeted nucleation in TEs

The restricted positioning of nucleation suggests a targeted mechanism for H3K9me3 deposition. Although one of the key functions of heterochromatin is to induce transcriptional silencing at TEs, a low level of nascent transcription has been implicated in the initial formation of heterochromatin in several organisms (Allshire and Madhani 2018). Transcription of satellites during embryogenesis has been shown to be required for the formation of heterochromatin in mouse and Drosophila (Probst *et al*. 2010; Casanova *et al*. 2013; Mills *et al*. 2019), and satellite transcription and/or RNAs are necessary for centromere and heterochromatin functions in humans, mice and flies (Rošić *et al*. 2014; Johnson *et al*. 2017; Shirai *et al*. 2017; Velazquez Camacho *et al*. 2017; McNulty *et al*. 2017). Chromatin-associated transcripts may recruit silencing factors to their genomic location (Holoch and Moazed 2015), and co-transcriptional silencing by Piwi-piRNA complexes has led to a model where piRNAs provide sequence specificity for PIWI to target nascent TE transcripts and guide co-transcriptional heterochromatin formation at TEs (Holoch and Moazed 2015). In Drosophila, the piRNA pathway is thought to be primarily active in gonadal tissue, yet mutations of key factors in the pathway, including Piwi, affect heterochromatin formation in somatic tissues (Pal-Bhadra *et al*. 2004; Gu and Elgin 2013). Both piRNAs and PIWI protein are maternally loaded into the egg, consistent with maternal piRNA/PIWI complexes guiding initial heterochromatin establishment in the early embryo, which is later maintained independently of piRNAs (Pal-Bhadra *et al*. 2004; Gu and Elgin 2013).

Supporting the requirement of nascent transcription for heterochromatin formation, we find that TEs showing early nucleation have significantly higher zygotic expression than the rest during early development (between stages 2 and 4 for TEs with stage3 peaks, or stages 4 and 5 for TEs with stage 4 peaks). In addition, we find that early nucleating TEs are associated with high abundance of both sense and anti-sense maternally deposited piRNAs. Early zygotic transcription during embryogenesis and high maternal piRNA counts are consistent with this model of co-transcriptional silencing of TEs, and highlight the importance of small RNAs in establishing heterochromatin in flies.

However, it is unclear whether piRNAs can account for the high specificity of early heterochromatin deposition at individual TEs. We find that TEs have specific regions that are targeted first for heterochromatinization, and the positions of abundant piRNA read mapping show little correspondence to the positions of H3K9me3 nucleation along TEs. Our observation that early nucleating TEs also show early zygotic transcription raises the possibility that the act of transcription at these TEs may provide specificity; indeed, extensive chromatin remodeling occur at promoters during zygotic genome activation (Schulz and Harrison 2019). Supporting this, the strong H3K9me3 enrichment (and large number of peaks) around the 5’ LTR of TRAM and other retrotransposons are consistent with a model where histone methyltransferases are recruited to the promoter of the TE insertions perhaps by transcription initiation. However, this does not appear to be a general rule across early nucleating TEs, as elements like R1-6 and LOA-2 have early nucleation sites close to the 3’ or near the middle of the insertions, respectively. These discrepancies may suggest that alternative targeting mechanisms are involved in early heterochromatin deposition in flies. KRAB-zinc-fingers are confined to mammals, but an analogous protein family exists in flies: ZAD-zinc-fingers (ZAD-ZNF’s). ZAD-ZNF’s are a poorly characterized gene family that has dramatically expanded within insect lineages (Chung et al. 2007)(Kasinathan *et al*. 2020). Approximately half of all ZAD-ZNF genes in *D. melanogaster* are highly expressed in ovaries and early embryos (Chung *et al*. 2002) and several interact with heterochromatin (Kasinathan *et al*. 2020), (Swenson *et al*. 2016). The ZAD-ZNF gene repertoire evolves rapidly across Drosophila species, which has been suggested to be driven by rapid alterations in heterochromatin across flies (Kasinathan *et al*. 2020). While it remains unclear how these genes localize to heterochromatin, it raises the possibility that maternally deposited DNA-binding factors can recruit histonemethyltransferases for H3K9me3 deposition in flies. The localized nature of the early nucleating sites is suggestive of a targeting mechanism such as motif recognition by DNA-binding proteins. Future research in Drosophila will show whether specific sequences on TEs are targeted by maternally deposited DNA-binding proteins that act in concert with the piRNA pathway to causes site-specific H3K9me3 deposition at recently active TEs.

### Early and robust silencing against highly active TEs

Regardless of the targeting mechanism of early H3K9me3 nucleation, early nucleating TEs are under robust silencing. They are disproportionately associated with high piRNA abundance, and early but low zygotic transcription. They also appear sufficiently silenced during oogenesis with the lowest maternal transcript abundance, and their expression remains significantly lower than the remaining TEs as zygotic transcription ramps up. What, then, sets these TEs apart and why are they more robustly silenced compared to others? Interestingly, we find that these TE are significantly more abundant with higher copy numbers and insertions across the genome. Elevated copy number on the neo-Y chromosome further indicates that these TEs had high transposition activity within the past 1.5 million years since the formation of the neo-Y. Altogether, these results reveal that the early nucleating TEs have high transposition potential and strong genomic silencing in the form of piRNA defense and targeted nucleation may have evolved to maintain genome integrity. Indeed, many components of the piRNA machinery and heterochromatin/TE regulating proteins are rapidly evolving (Blumenstiel *et al*. 2016). In the ongoing arms race between proliferation of selfish repeats and genome defense, targeted early H3K9me3 nucleation may provide complementary or redundant mechanisms to piRNA-mediated transcriptional silencing.

## Materials & Methods

### Embryo collection and staging

*D. miranda* adults were allowed to lay for 1 hour in an embryo collection cage at 18°C to pre-clear any old embryos that females may be holding. After pre-clearing, the embryos were aged to their appropriate developmental stage. Embryos were then dechorionated with 50% bleach solution for 1 min and identified according to its morphological features under a light microscope. Embryo developmental time/stage: stage3 (210min, polar buds at the posterior pole of the embryo), early stage4 (240min, thin “halo” appears as the syncytial blastoderm forms), late stage4 (270min, thicker “halo” compared with early stage4 and no formation of cell membranes), stage5 (300min, visible cell membranes as cellularization progresses), stage7 (420min, cephalic furrow).

### Single embryo chromatin immunoprecipitation and library preparation with spike-in

The H3K9me3 ULI-nChIP preparation and pull down were done according to a modified version of (Brind’Amour *et al*. 2015). In short, *D. miranda* embryos single embryos were homogenized into a cell suspension using a 200ul pipette tip in 20ul of Nuclei EZ lysis buffer (Sigma-Aldrich Catalog No. N3408). To fragment the chromatin, 2 U/ul of MNase to were added into each nuclei preparations for 7.5 minutes at 21°C; reaction was terminated by adding XXM EDTA. Simultaneously, *D. melanogaster* stage 7 embryos were prepared the same way and 5ul was added into each *D. miranda* samples for spike-in. Then the digested nuclei preps were diluted to 400ul with ChIP buffer (20 mM Tris-HCl pH 8.0, 2 mM EDTA, 150 mM NaCl, 0.1% Triton X-100, 1x Protease inhibitor cocktail) and split into a 40ul aliquat and 360ul aliquat for input and ChIP, respectively. Prior to the ChIP pull down, antibody against H3K9me3, (Diagenode Cat No. C15410056), H4 (Abcam ab1791), H3K4me1 (Diagenode Cat. No. C15410194), H4K16ac (Millipore Cat. No. 07-329) were incubated with Dynabeads (ThermoFisher Cat No. 10002D) for 3hr to make the antibody-bead complex, followed by overnight incubation with the chromatin preparations. DNA from ChIP and input samples were isolated with Phenol-chloroform using MaXtract High Density tubes (Qiagen Catalog No. 129046) to maximize sample retrieval, followed by ethanol precipitation and resuspension in 30ul of nuclease-free water. The resulting DNA extractions were further purified to remove excess salt using Ampure XP beads at 1.8:1 beads-to-sample ratio (54 ul) and eluted in 10ul of nuclease-free water. Library preparations were done using the SMARTer Thruplex DNA-seq kit from Takara (cat# R400674). Library concentration was determined using Qubit (Thermofisher) and the library quality was determined using the Bioanalyzer by the Functional Genomics Lab at UC Berkeley.

### ChIP Sequencing and data processing

All libraries were (100bp) paired-end sequenced with the Hi-Seq 4000 at the Vincent J. Coates Genomics Sequencing Lab at UC Berkeley. To differentiate between spike-in and sample from each ChIP-seq library, raw paired-end reads are aligned to a reference file that contains both the *D. miranda* genome (r2.1) and the *D. melanogaster* genome (r6.12) using bwa mem (Li and Durbin 2009) on default settings. Since bwa mem, by default, only reports the highest quality alignment, each read will align to one of the two genomes only once. When there are more than one alignment of equal mapping quality, the read will be randomly assigned to one of the targets. Using a custom Perl script, reads are then differentiated based on which of the two genomes they align to, and ambiguous alignments are discarded. For read mapping statistics, see **Supplementary Table 1**. Reads are then sorted using samtools (Li *et al*. 2009) and the coverage per site is determined with bedtools genomecoverge (Quinlan and Hall 2010) (-d -ibam options).

Sex of embryos were inferred from the ratio of the median autosomal coverage: median X coverage of the inputs (**Supplementary Table 1**). Median coverages were inferred from distribution of average coverages in 1kb sliding windows.

### Sample and enrichment normalization

For normalization without spike-in, we first took the average coverage of the ChIP and input samples in 1kb windows, and then determined the median of all the autosomal windows to obtain the median autosomal coverage. The per-site coverages in the ChIP and input samples were then divided by their respective median autosomal coverage, thus normalizing for library size. The per-site enrichment is then estimated as:

Enrichment = (ChIP/ChIP median + 0.01) / (input/ChIP median + 0.01),

with, 0.01 being a small pseudo-count.

For normalization with spike-in, we first generated a reference spike-in enrichment profile by using the same procedure as above for all spike-in ChIPs and spike-in inputs in 1kb windows, and then took the average log_2_ enrichment across all spike-ins, generating a reference. This reference is meant to be a standard to which all other spike-ins will be normalized, and the extent of normalization in the spike-ins will be applied to the actual samples. To do so, we used a method akin to quantile normalization, whereby the distribution of the spike-in sample is matched with that of the reference. However, instead of matching the distributions identically as with typical quantile normalization, we binned the enrichment values into quantiles of 0.1, allowing us to match the quantiles by bins. For example, if the 98.1^th^ quantile has a log2 enrichment of 3.2 and 4.0 in the spike-in sample and reference, respectively, all points of the spike-in within the quantile will be increased by 0.8. Since the spike-in and the actual sample should have the same pull-down efficiency, this then allows us to determine the extent of normalization to apply to each log_2_ enrichment in the actual latter; for example, points with log_2_ enrichment of 3.2 will similarly be elevated by 0.8 in the sample.

### ChIP-seq peak calling

We used macs2 (Zhang *et al*. 2008) to call H3K9me3 enrichment peaks. For each stage containing multiple replicates, we called peaks using sorted bam files with the spike-in reads removed, where the spike-in reads are removed with the command below:

macs2 callpeak -t replicate1.chip.sort.bam replicate2.chip.sort.bam … -c replicate1.input.sort.bam replicate2.input.sort.bam … -n output_name -g 1.8e8 -call-summits

For peak calling in individual replicates, only the chip and corresponding input bam files are used for the -t and -c parameters. Peaks in different replicates that are less than 100bp away from each other are deemed the same.

### Enrichment analysis around peaks

For each developmental stage, we averaged the H3K9me3 enrichment per site (or bin) across the replicates to generate a representative enrichment. To determine the developmental progression of enrichment around peaks, we averaged the representative enrichment at 100bp upstream and downstream of the summit of the peak (reported by macs2) for each stage. Regions are deemed enriched if log_2_(enrichment) > 0.5. Enrichment on the neo-Y is averaged from only male embryos. For the enrichment heatmaps, we took the per-base enrichment 1000bp upstream and downstream of the summits and sorted the peaks from highest enrichment to lowest enrichment. We then plotted each base as a dot using colors that scale with the enrichment with a custom R script. Colors are generated using the R package RColorBrewer. Heatmaps around peaks of ATAC-seq, RNA-seq, and DNA-seq are also generating using this method.

### Repeat annotations, contents, and overlap with peaks

We used Repeatmasker (Smith *et al.)* with a TE index specific to the *obscura* group (the Drosophila lineage to which *D. miranda* belongs) from (Hill and Betancourt 2018). Simple repeats were identified using the simple repeat setting in Repeatmasker which lists them and can be identified in the annotation file (.gff) as (motif)n where the “motif” is the repeating unit (e.g. (AGAT)n). To determine the %repeat content, we used the repeat masker annotation and determined the number of bases annotated by either a TE or a simple repeat in nonoverlapping sliding windows. To determine overlap between annotated repeats and H3K9me3 peaks, we used bedtools intersect -a annotatedrepeats.gff -b macs2.peaks -wa -wb. Distance between repeats and peaks is inferred using the R package IRange (Lawrence *et al*. 2013)

### H3K9me3 enrichment at full length and truncated annotated TE insertions

The Repeatmasker annotation (gff) provides the positions of the TE insertions in the genome as well as well the positions matching the TE consensus sequence; for each TE entry, these coordinates are used to create a table of enrichment values where each row is an annotated insertion in the genome and each column is a position of the consensus from the repeat index. Because the LTR and internal sequences of LTR retrotransposons are separate entries in the repeat index, we identified full length insertions of TRAM in the genome by identifying internal sequences flanked by LTRs requiring that all three features must have the same direction (all forward/reverse) and are within 10bp of each other. To identify 5’ and 3’ truncated insertions, we identified internal sequences where the respective LTRs are either missing, or greater than 5kb away. For statistical comparisons, we averaged the H3K9me3 enrichment across the last 1kb of the element of each full length and truncated insert, and compared the distribution of averaged enrichment between full length and 5’ insertions. The opposite was done for 3’ truncated insertions, where we averaged across the first 1kb.

### DNA-seq and RNA-seq data analysis

Sexed DNA-seq and RNA-seq data are from (Mahajan *et al*. 2018) and (Lott *et al*. 2014). Raw reads were aligned using bwa mem on default settings to either the *D. miranda* genome or the Repeat index (Hill and Betancourt 2018), and sorted with samtools. After samtools sort, the per-base coverage is determined with bedtools genomeCoverageBed and then normalized by the number of mapped reads in the library. For copy-number-scaled TE transcript abundance, we tallied the RNA-seq and DNA-seq read count mapping to each TE entry in the repeat library; the former was normalized by the median read count at autosomal genes and the latter normalized by the median autosomal coverage. The RNA-seq read counts were then divided by the DNA-seq read counts for each TE.

### Embryo collection for piRNA sequencing and piRNA isolation

Flies were allowed to lay on molasses plates with yeast paste for one hour and embryos were immediately flash frozen in liquid nitrogen and stored at −80°C. We used Trizol (Invitrogen) and GlycoBlue (Invitrogen) to extract and isolate total RNA. piRNA is isolated using NaIO_4_ reaction and beta elimination with a protocol modified from (Kirino and Mourelatos 2007; Ohara *et al*. 2007; Cao *et al*. 2009; Simon *et al*. 2011; Abe *et al*. 2014; Gebert *et al*. 2015). We started with 20 ug total RNA in 13.5 ul water and added 4 ul 5x borate buffer (148 mM borax, 148 mM boric acid, pH 8.6) and 2.5 ul freshly dissolved 200mM sodium periodate. After 10 minutes of incubation at room temperature, the reaction is quenched with 2 ul gylcerol for an additional 10 minutes. We poked holes in the tops of the tubes with sterile needles, then vacuum dried the reaction for 1 hour. We then added 50 ul 1x borax buffer (30 mM borax, 40 mM boric acid, 50 mM NaOH, pH 9.5, and 200 mM sodium periodate) and incubated the samples f**or** 90 minutes at 45°C. For RNA precipiration, we added 1.33 ul GlycoBlue (Invitrogen) and 200 ul ice cold 100% EtOH and place the samples in −80°C overnight. We centrifuged down the samples at 4°C, maximum speed, for 15 minutes, removed the supernatant, and let the RNA pellet dry, and resuspended the RNA in 20 ul H_2_O.

20 ul of RNA prepared as described above was resolved on a 15% TBE-urea gel (Invitrogen). Custom 18-nt and 30-nt ladders were used to size select 19-29 nt long RNA, which was purified and used as input for library preparation using the Illumina TruSeq Small RNA Library Preparation Kit to ligate adapters, reverse transcribe and amplify libraries, and purify the cDNA constructs. The protocol included an additional size selection step after library preparation on a 6% TBE gel (Invitrogen). 50bp single-end sequencing of our samples was performed on an Illumina HiSeq 4000 at the Vincent J. Coates Genomic Sequencing Laboratory at UC Berkeley.

### piRNA-seq data analysis

Adapters were trimmed from the reads using trim_galore (http://www.bioinformatics.babraham.ac.uk/projects/trim_galore/). We mapped the reads to the *D. miranda* genome and the repeat index using the small RNA-specific aligner ShortStack (v3.8.5) (Johnson *et al*. 2016). Using samtools view we then extracted reads that are 23-29bp in length. We then used bedtools genomeCoverage to obtain piRNA coverage at each position in the genome or repeat index.

### ATAC-seq sample preparation

Single staged embryos were collected as above. ATAC-seq preparation was done according to a modified protocol of (Buenrostro *et al*. 2015). Embryos were homogenized into a cell suspension using a 200ul pipette tip in 20ul of Nuclei EZ lysis buffer (Sigma-Aldrich Catalog No. N3408). The supernatant was removed after centrifugation at 4°C. The nuclei were the resuspended in the transposition buffer (2x reaction buffer, Illumina Cat #FC-121-1030, Nextera Tn5 Transposase, Illumina Cat #FC-121-1030) and PCR amplified (KAPA HiFi HotStart ReadyMix, cat# KK2600) for 12 cycles. PCR products were cleaned up with AMPure XP beads (cat# A63881) at 1:1 concentration. Samples were sequenced with the Novaseq 6000 S1 with 100bp pair-end reads at the Vincent J. Coates Genomic Sequencing Lab at UC Berkeley.

### ATAC-seq data analysis

Pair-end reads are adapter trimmed with trimgalore and aligned to the reference genome with bwa mem on default settings. After sorting with samtools, we used bedtools genomeCoverageBed with the -pc flag for fragment count instead of read count, generating the per base fragment coverage. The fragment coverage for each sample is then normalized by the median autosomal coverage. The sex of the embryos is then determined based on the coverage on the sex chromosomes; I.e. males will have ~0.5x coverage on the X and Y chromosomes. Since our analyses focused on repetitive heterochromatin where read coverage deviates highly due to copy number differences, we controlled for copy number by dividing the ATAC-seq fragment counts by fragment counts of sex-matched DNA-seq samples, generating ATAC enrichment. This also simultaneously controls for the coverage differences of sex chromosomes between males and females.

## Data availability

All ChIP-seq and ATAC-seq data generated have been deposited on Genebank under BioProject PRJNA601450 (currently under embargo). Intermediate files, including ChIP enrichment files and peak calls, are uploaded on Dryad. R and perl scripts for spike in normalization, generating enrichment around peaks, and enrichment heatmaps are available on KW’s github page (https://github.com/weikevinhc/heterochromatin.git)

## Notes

### Competing Interest Statement

The authors have declared no competing interest.

